# DOCK2-deficiency causes defects in anti-viral T cell responses and poor control of herpes simplex virus infection

**DOI:** 10.1101/2023.08.02.551154

**Authors:** Katrina L. Randall, Inge E.A. Flesch, Yan Mei, Lisa A. Miosge, Racheal Aye, Zhijia Yu, Heather Domaschenz, Natasha A. Hollett, Tiffany A. Russell, Tijana Stefanovic, Yik Chun Wong, Christopher C. Goodnow, Edward M. Bertram, Anselm Enders, David C. Tscharke

**Affiliations:** Division of Immunology and Infectious Diseases, John Curtin School of Medical Research, Australian National University, Canberra, ACT 2601; School of Medicine and Psychology, Australian National University, Canberra ACT 2600; Garvan Institute of Medical Research, University of New South Wales, Darlinghurst, NSW 2010, Australia

**Keywords:** Dedicator of cytokinesis 2, DOCK2, herpes simplex virus, T cell activation, viral control

## Abstract

The expanding number of rare immunodeficiency syndromes offers an opportunity to understand key genes that support immune defence against infectious diseases. However, patients with these diseases are by definition rare. In addition, any analysis is complicated by treatments and co-morbid infections requiring the use of mouse models for detailed investigations. Here we develop a mouse model of DOCK2 immunodeficiency and demonstrate that these mice have delayed clearance of herpes simplex virus type 1 (HSV-1) infections. Further, we found that they have a critical, cell intrinsic role of DOCK2 in the clonal expansion of anti-viral CD8^+^ T cells despite normal early activation of these cells. Finally, while the major deficiency is in clonal expansion, the ability of primed and expanded DOCK2-deficient CD8^+^ T cells to protect against HSV-1-infection is also compromised. These results provide a contributing cause for the frequent and devastating viral infections seen in DOCK2-deficient patients and improve our understanding of anti-viral CD8^+^ T cell immunity.

## Introduction

The management of infectious diseases in patients with primary immunodeficiency is a significant clinical problem. At the same time, the expanding catalogue of primary immunodeficiencies is revealing not only new roles for mammalian genes in immunity, but also an appreciation that many gene defects lead to unique susceptibility to infectious diseases [1]. DOCK2 immunodeficiency is a disease that leads to severe immunocompromise, being fatal in two of the original five cases described, and requiring bone marrow transplantation in the other cases [2]. Patients with mutations in *DOCK2* present with combined immunodeficiency with early onset invasive bacterial and viral infections [2]. Typical infections found in the published DOCK2 patients include invasive viral infections including varicella, mumps, cytomegalovirus and adenovirus, as well as bacterial infections and a likely case of Pneumocystic jirovicii [2-4]

DOCK2 is a member of the DOCK family of guanine nucleotide exchange factors (GEFs) and has been previously shown in mice to be a GEF for RAC1 [5, 6]. DOCK2 is predominantly expressed in hematopoietic cells, particularly the cells of the immune system [7]. Previous studies using a DOCK2 knockout mouse have shown that loss of DOCK2 is associated with severe peripheral lymphopenia and lymphoid follicle hypoplasia [7]. DOCK2 has been shown to be important for proper T cell synapse formation after activation by antigen and aids in the translocation of the T cell receptor (TCR) and lipid rafts into the synapse [5]. DOCK2 had also been shown to be important for integrin activation in response to chemokine signaling in B cells [8].

In the more severe syndromic immunodeficiencies, it can be especially difficult to dissect the ways in which a particular gene defect compromises control of a given pathogen. Multiple concurrent infections and medications can mask or exacerbate immune consequences of the defects in these patients. Therefore for a gene like *DOCK2*, with roles in multiple cell types, reductionist models are required. In this regard, mouse models are particularly useful. Indeed one study has shown that DOCK2 is important for protection against enteric infection with *Citrobacter rodentium*, with a role for this protein in preventing or reducing bacterial attachment to enterocytes being identified [9], as well as effects on macrophage migration [10]. However, such an innate mechanism seems unlikely to underlie a susceptibility to viral infections, nor does it articulate well with the known requirement for DOCK2 in lymphocytes.

Here we take advantage of a well described model of viral skin infection with herpes simplex virus (HSV) in mice with DOCK2 deficiency to examine this defect in the context of a viral infection. We show that DOCK2 deficient mice have a more severe disease after HSV infection, including greater lesion size and increased viral titres. This model was then extended to explore anti-viral CD8^+^ T cell function. This found a major cell-intrinsic defect in expansion of virus-specific CD8^+^ T cells and a lesser, but still significant deficiency in protective capacity. Consistent with these findings, the numbers of endogenous virus-specific CD8^+^ T cells were reduced in mice acutely infected with HSV. These data provide insight into the impact of DOCK2 deficiency on anti-viral CD8^+^ T cells.

## Methods

### Viruses and cell lines

HSV-1 strain KOS was kindly provided by F. Carbone (The University of Melbourne, Parkville, Victoria, Australia) and is referred to as HSV throughout. HSV.OVA is a recombinant of HSV-1 strain KOS expressing a fusion of enhanced green fluorescent protein and the epitopes SIINFEKL, TSYKFESV, SSIEFARL and has been described previously [11]. Viruses were grown and titrated by standard methods using BHK-21 for growth and BS-C-1 for titration, respectively. Immortalized cell lines BHK-21 and BS-C-1 were maintained in Dulbecco’s Modified Eagle medium (DMEM, Invitrogen) with 2 mM L-glutamine and 10% fetal bovine serum (FBS) (D10). Vero cells were grown in Minimal Essential Medium supplemented with 10% FBS, 2 mM L-glutamine, 5x10^-5^ M 2-mercaptoethanol (2-ME) and 5 mM HEPES (all Invitrogen).

### Mice

Specific pathogen-free female C57BL/6, C57BL/6.SJL (CD45.1) and C57BL/6 OT-I mice greater than 8 weeks of age were obtained from the Animal Resource Centre (Perth, Australia) and from the Australian Phenomics Facility (APF, Canberra, Australia). DOCK2*^E775X/E775X^* (ENSMUST00000093193) mice were generated by chemical mutagenesis using N-ethyl-N-nitrosourea (ENU) as previously published [12, 13]. ENU was given intraperitoneally (i.p.) to male C57Bl/6 mice three times at an interval of 1 week. All mice were housed, and experiments were done according to the relevant ethical requirements and under approvals from the ANU animal ethics and experimentation committee (A2011/01, A2013/37, A2014/62, A2016/45, A2017/54, A2020/01 and A2020/45) at the APF.

### HSV infections

Female mice >8 weeks of age were anesthetized by i.p. injection of Avertin (20 µl/g of body weight). The left flank of each mouse was shaved and depilated with Veet. HSV was diluted in PBS to 10^8^ PFU/ml and tattooed into a 0.5x0.5 cm area of skin above the tip of the spleen. Body weight and lesion progression were measured daily until the lesions had resolved. Lesion size was determined with the aid of a caliper to determine overall area and then the proportion of the area affected by the lesion was estimated and used to calculate a final size. In some experiments spleens were taken after seven days and cells analyzed for HSV-gB_498_-specific CD8^+^ T cells, or CD8^+^ T cells that make IFN_γ_ after stimulation with gB_498_ peptide, by flow cytometry (see below).

### Viral titer determination

Dorsal root ganglia (DRG) innervating the infected dermatome were removed at day 7 post infection. All DRG from one mouse were pooled into 1 ml of DMEM supplemented with 2% FBS and 4 mM L-glutamine (D2). Samples were homogenized, freeze-thawed three times and viral titers were determined using standard plaque assays on monolayers of confluent Vero cells and expressed as plaque forming units (pfu) per mouse [14].

### Activation of OT-I T cells in vitro for analysis and HSV protection

Splenocytes were prepared from D2EX and WT littermate OT-I mice. For in vitro analysis experiments, 2×10^6^ splenocytes were cultured with OVA_257_ peptide (SIINFEKL, concentrations as shown) in D10 supplemented with 5x10^-5^ M β-mercaptoethanol and 5 mM HEPES (T cell medium) for up to 40 hours before harvesting and flow cytometric staining for either CD69 or intracellular IRF4. For preparation of bulk cultures of OT-I T cells for transfer into mice, splenocytes were prepared as above, but cultures were started with 1×10^8^ splenocytes. One third of these were pulsed with 1×10^-7^ M OVA_257_ peptide in serum-free medium for 1 hour at 37°C on a rocking platform before washing and recombining with the other cells. Cultures proceeded in T cell medium, further supplemented with recombinant IL-2. Cultures of D2EX OT-I failed unless supplemented with higher amounts of IL-2 and we determined empirically that using 6 ng/ml for D2EX OT-I produced cultures of cells similar to WT OT-I in 3 ng/ml, so these differing amounts of cytokine were used. After 4 days cultures were enriched for CD8^+^ T cells using a MACS CD8a^+^ T Cell (untouched) Isolation Kit (# 130-095-236) according to manufacturer’s instructions. 5×10^6^ purified cells (typical purity <90% CD8^+^) were transferred into female WT mice (>8 weeks old) via i.v. injection in a total volume of 200 µl PBS. Control mice received 200 µl PBS. Twenty-four hours later, mice were tattoo-infected with HSV.OVA (as above).

### Activation and expansion of naïve OT-I CD8^+^ T cells by HSV infection in vivo

Splenocytes were prepared from D2EX and WT littermate CD45.1^+^ OT-I mice and enriched for CD8^+^ T cells using a MACS CD8a^+^ T Cell (untouched) Isolation Kit (# 130-095-236) according to manufacturers instructions. After purification cells were typically ∼90% CD8+, Vα2^+^. 1×10^4^ of these cells were injected i.v. into female CD45.2^+^ recipient mice (>8 weeks old) that were then infected on the flank with HSV.OVA 24 hours later (as above). Seven days after infection, mice were culled and numbers of OT-I cells in the spleen and/or DRG identified as CD8^+^, CD45.1^+^, Vα2^+^ events by flow cytometry.

### Flow cytometry

Blood was collected from the retroorbital veins using EDTA as anti-coagulant. Single-cell suspensions from organs were prepared by mashing organs through a 70µm cell strainer (BD) followed by antibody staining as described previously [15]. Erythrocytes in blood and spleen samples were lysed using ammonium chloride lysis buffer before antibody staining. Peripheral blood screen: AlexaFluor700 (AF700)-conjugated anti-CD4 (BD, RM4-5), peridin-chlorophyll-protein complex (PerCP)-Cyanine (Cy) 5.5 conjugated anti-B220, Pacific Blue (PB)-conjugated anti-CD44, allophycocyanin (APC)-Cy7-conjugated anti-CD3, APC-conjugated anti-NK1.1 (PK136, BD), fluorescein isothiocyanate (FITC)-conjugated anti-IgM (, phycoerythin (PE)-Cy7 conjugated anti-KLRG1, and PE-conjugated anti-IgD. Thymic and splenic surface stains (eBioscience unless otherwise stated): AF700 conjugated anti-CD4 (BD, RM4-5), Brilliant Ultraviolet (BUV) 395-conjugated anti-CD8 (BD, 53-6.7), APC-conjugated anti-CD5 (53-7.3), PE conjugated anti-CD25 (PC61.5), PerCP Cy 5.5-conjugated anti-CD3 (BioLegend, 17A2), PE-conjugated anti-CD3 (BD, 145-2C11), Brilliant Violet (BV) 605 conjugated anti-CD62L (Biolegend, MEL-14), PB-conjugated anti-CD44 (BioLegend, IM7), APC-Cy7-conjugated live/dead stain, FITC conjugated anti-TCR-β (H57-597, eBioscience), efluoro conjugated live/dead stain, biotin-conjugated anti-CD93 (AA4.1), PE-Cy7-conjugated IgM (II/41), FITC conjugated anti-IgD (11-2c (22-26)), PB-conjugated anti-CD23 (B3B4), BUV737-conjugated anti-CD21/35 (BD, 7G6), AF700-conjugated anti-B220 (RA3-6B2) and BUV395-conjugated anti-CD19 (BD, 1D3).

All B cell stains included Fc block (BD, 2.4G2), either as a 30 minutes pre-incubation or together with biotinylated or fluorescently labelled antibodies. Biotin staining was followed by addition of BV605-conjugated streptavidin (BioLegend). For intracellular staining of FOXP3, the eBioscience Foxp3 / Transcription Factor Staining buffer set (00-5523-00) was used according to the manufacturer’s instruction using FITC-conjugated anti-FoxP3 antibodies (FJK-16S). Detection of NKT cells using CD1d monomers loaded with α-GalCer (produced by the NIH tetramer facility) was as previously described [16].

3) For CD8^+^ cells and HSV-specific CD8^+^ T cells in infected mice, Surface stain panel H-2Kb/SSIEFARL dextramer (Immudex), anti-CD8 (clone 53-6.7; BioLegend) and in some cases anti-CD62L (MEL-14, BioLegend) and intracellular staining with anti-GzmB (GB11, BioLegend). 4) After stimulation with gB_498_ peptide (SSIEFARL) for 4 hours in the presence of brefeldin A, anti-CD8 (as above) and anti-IFNγ (XMG1.2, BioLegend), stained intracellularly[17]. 5) For OT-I cells tested prior to transfer or from mice after transfer and infection, anti-CD8 (as above), anti-CD45.1 (A20, Biolegend) and anti-TCRVα2 (B20.1, BioLegend). 6) For OT-I cells stimulated in vitro, anti-CD8 (as above) and anti-CD69 (HI.2F3, BD Bioscience) or anti-IRF4 (3e4, eBioscience) stained intracellularly using a Foxp3 / Transcription Factor Staining Buffer Set (cat# 00-5523-00, eBioscience). Samples were acquired on a LSR II flow cytometer and analysis was done using Flowjo software (Tree Star Inc.). Statistical analysis was done using GraphPad Prism.

## Results

### Novel DOCK2 mutant mouse strains generated by ENU mutagenesis

As part of an ENU-mutagenesis project to provide mouse models for human disease [12], 3 different mouse strains with premature stop codons in *DOCK2* were discovered due to T cell lymphopenia in the blood as shown in Figure 1A. The position of the mutations in the DOCK2 protein are shown in Supplementary Figure 1A.

**Figure 1:**
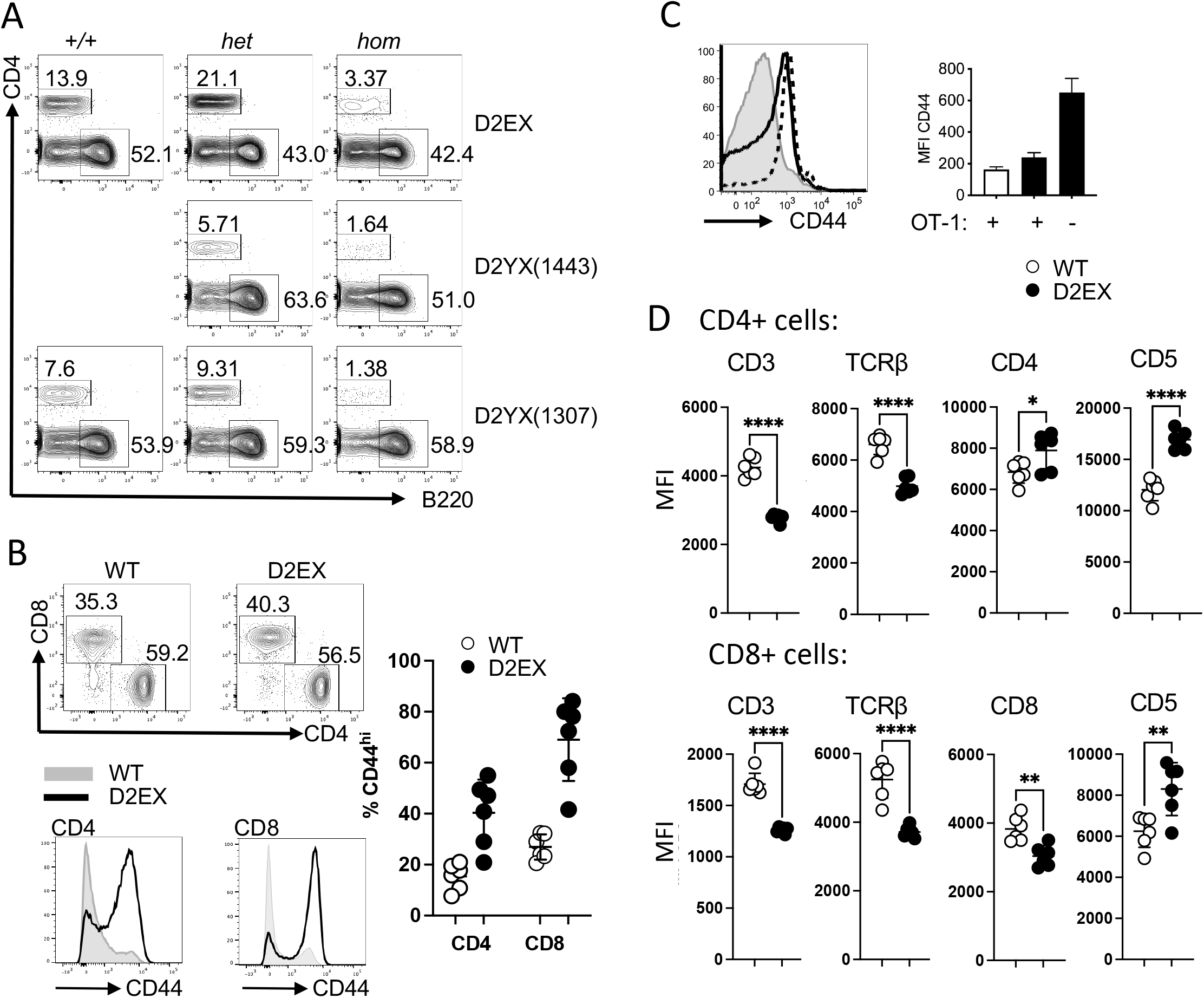
CD4+ T cell lymphopenia in the absence of DOCK2. A)Representative flow cytometry plots (pre-gated on lymphocytes) from mice carrying two copies (*hom*), one copy (*het*) or no copies (+/+) of the listed amino acid change in DOCK2. B) Representative flow cytometry plots (pre-gated on CD3+ lymphocytes) from wild type and mutant mice (top panel) and quantitation of proportion of CD44hi CD4+ and CD8+ T cells from the two groups (right panel). Representative histograms showing the CD44 staining of CD4+ and CD8+ T cells from wild type (grey) and mutant mice (black line). Representative of at least 3 separate experiments. C) Effect of limiting TCR repertoire on the expansion of CD44+ lymphocytes. Representative histogram of CD44 expression for wild-type mice with OT-I (grey), D2EX mice with OT-I expression (dotted) and D2X mice without OT-I (black line), and quantitation of MFI for these groups of mice - wild-type mice with OT-I (white bar), D2EX mice with OT-I expression (black bar) and D2X mice without OT-I (black bar) with absence/presence of OT-I noted on x axis. D) Relative expression of the listed markers on wild type and mutant CD4+ (upper panel) and CD8+ (lower panel) T cells. Unpaired t-test. * p<0.05, ** p<0.005, **** p < 0.0001.

### Characterization of the DOCK2 E775X strain

One of the strains strain carrying the E775X mutation due to a G to T point mutation at position 2392 in cDNA (ENSMUST00000093193) was selected for further analysis. Homozygous mice carrying this mutation (i.e. DOCK2*^E775X/E775X^*) are referred to hereafter as D2EX for brevity. This mouse strain recapitulates the already published features of DOCK2 mutation in mice, with marked T cell lymphopenia [7], in the blood of mice homozygous for the E775X mutation despite overall normal numbers of leucocytes (Figure 1A and Supplementary Figure 1B), absent marginal zone B cells [7] and decreased NKT cells in the thymus [18](Supplementary Fig 1C) with some increase of monocytes and eosinophils, and normal number of lymphocytes. We also detected elevated levels of IgE with aging in these mice (data not shown).

Closer analysis of T cell subsets in the spleen of these mice shows that the majority of the T cells (both CD4 and CD8) have an activated CD44 high phenotype (Figure 1B). This activation phenotype was partially ameliorated in mice with a transgenic T cell receptor (OT-I mice) with the mean fluorescence intensity of the whole population for CD44 decreasing on CD8+ transgenic cells but it is not completely normalized (Figure 1C). Interestingly, we found that the average expression of CD3 and TCRβ were decreased on mutant T cells. Furthermore, expression of the CD8 co-receptor on CD8+ T cells was decreased but expression of CD4 was increased on mutant CD4^+^ T cells. In line with a dysregulated TCR signaling in mutant T cells, we find that CD5 expression is increased on both CD4 and CD8 T cells in the spleen (Figure 1D).

We also enumerated FoxP3^+^ Tregs in the spleen and found that both their percentage and numbers were increased (Figure 2A). Despite the peripheral T cell lymphopenia, thymic T cell subsets in DOCK2 mutant mice were comparable to WT littermates (Figure 2 B), however thymic NKT cells were reduced (Supplementary Figure 1D).

**Figure 2:**
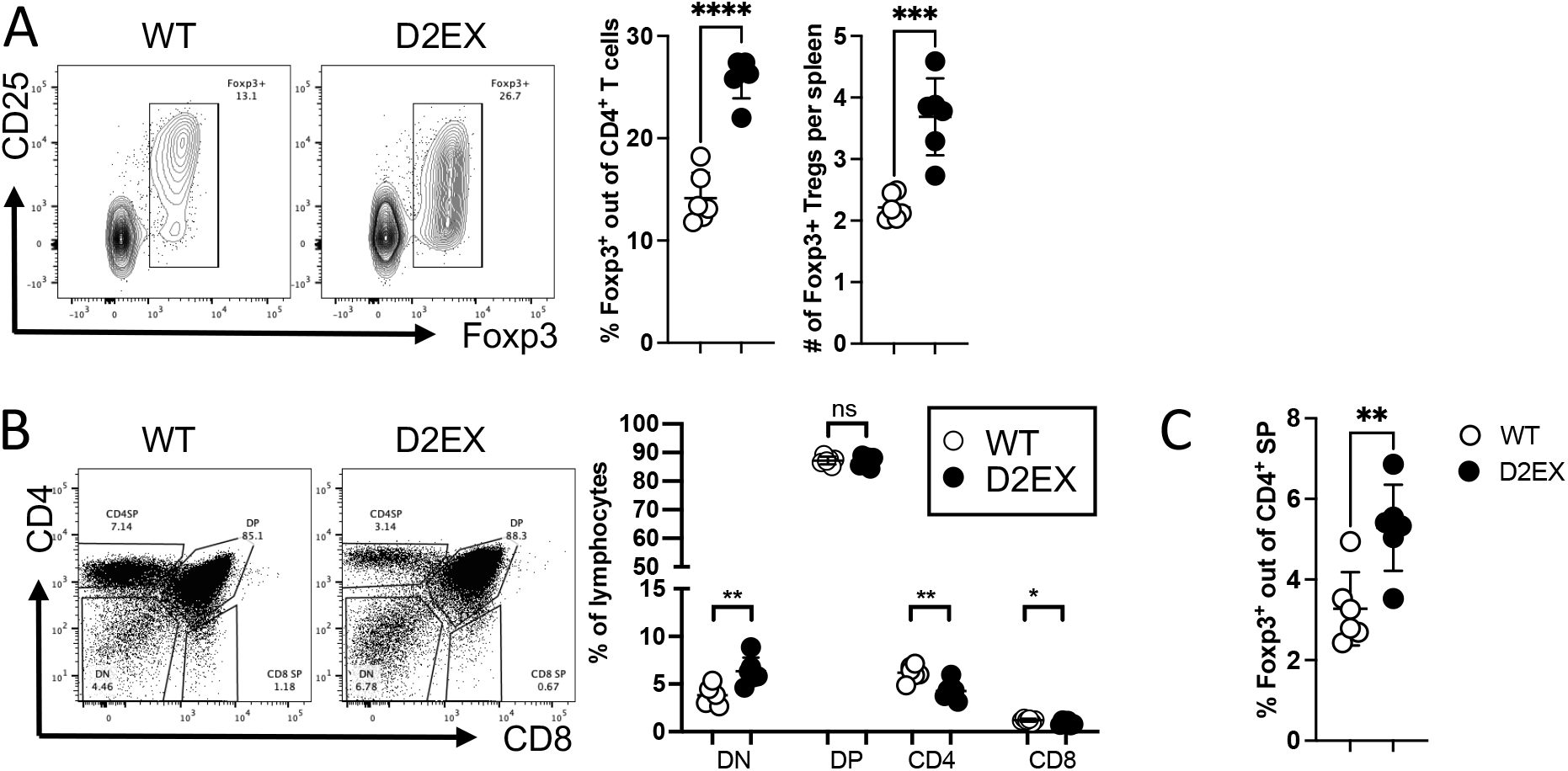
Increased formation of Foxp3^+^ Tregs in the absence of DOCK2. A) Naïve mice were analyzed by flow cytometry for the % and number of splenic Foxp3+ Tregs. B) Thymic T cell development was analyzed in naïve mice C) Thymic Foxp3+ cells were increased as a percentage of CD4SP T cells. Unpaired t-test. * p<0.05, ** p<0.005, **** p < 0.0001. Data representative of 3 independent experiments.

### D2EX-mutant mice loose significantly more weight and develop bigger lesions after skin infection with HSV-1

Cohorts of D2EX and wild-type C57BL/6 mice were inoculated with HSV.KOS in the flank. In this model, productive infections begin in the skin, but the virus then rapidly invades the peripheral nervous system, where further infection ensues in primary sensory neurons. Following spread in the nervous system, virus then emerges to other cutaneous sites throughout the infected dermatome producing a rash that is reminiscent of herpes zoster [19, 20]. Infection with this strain of HSV is very rarely lethal in mice and lesion size and weight loss can be assessed daily as clinical signs that indicate the severity of infection[14] After infection of WT and D2EX mice, weight and lesion progression were measured daily until the lesions had resolved and weight had reached the starting point of 100%. In both groups of mice, weight dropped sharply on days 1 and 2 after infection, but thereafter WT mice gained significantly more weight than D2EX mutant mice from day 10 to 20 post infection (Figure 3A). Further, D2EX mice developed significantly larger lesions from day 7 to 10 and while lesions were resolved by day 10 in wild-type mice, in D2EX mice lesions did not resolve for a further three days (Figure 3B).

**Figure 3:**
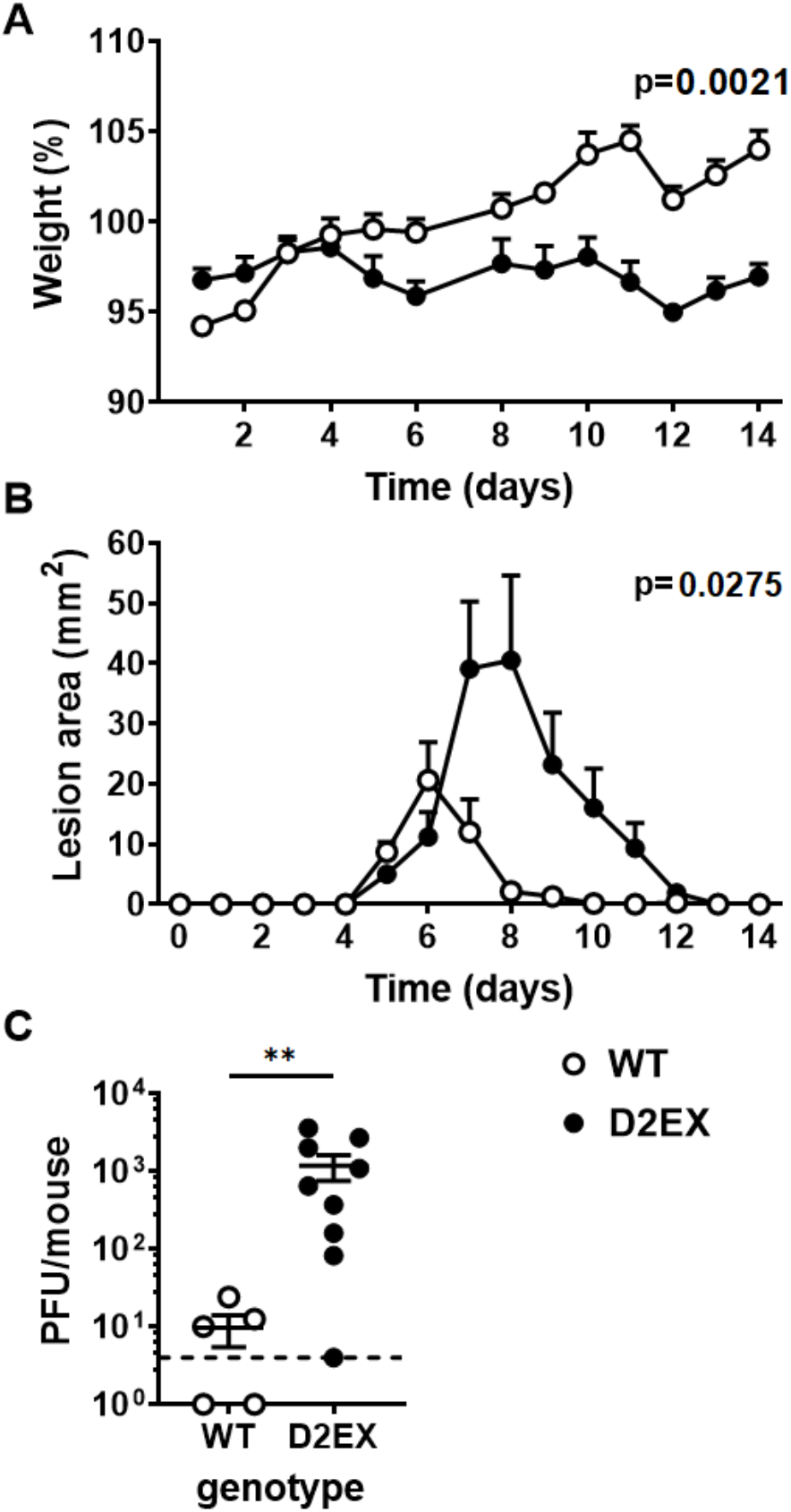
Delayed clearance of HSV in the absence of DOCK2. Mice were infected with HSV on the flank and pathogenesis (A and B) and viral loads (C) measured. A) Weights and lesion sizes of groups of 6 WT and 7 D2EX mice were monitored for 14 days. Differences between the strains were determined by two-way ANOVA, with significant p-values noted in the top right of graphs. C) Loads of infectious virus in the DRG of mice were measured by plaque assay 7 days after infection and the difference in means was tested using a t-test (**p<0.01). The experiment in A and B is representative of 3 independent repeats. C shows data combined from two independent experiments.

### Viral titers in DRG are higher in D2EX mice at day 7 post skin infection

The difference in pathogenesis suggested that the main impact of the defect in D2EX mice was to delay the clearance of infection that typically occurs with the effective deployment of activated T cells between days 5 and 8 after infection [19]. To test this, groups of D2EX and WT mice were infected and levels of HSV in DRG were quantified seven days later. In WT mice, two of five mice had already cleared virus to below the limit of detection and the average titre for the group was 10 pfu per mouse. By contrast only one of nine D2EX mice had undetectable virus and the average was 100-fold higher than seen in the WT mice (Figure 3C).

### DOCK2 has a cell-intrinsic role in mounting anti-viral CD8^+^ T cell responses

HSV infection of mice has provided an excellent model for interrogating CD8^+^ T cell priming, expansion and function [21-25] and is relevant to human infection [26, 27]. Therefore we bred D2EX mice to the OT-I T cell receptor (TCR)-transgenic mouse line to examine the activation and expansion of CD8^+^ T cells in response to infection with HSV.OVA, which expresses the SIINFEKL epitope recognised by the OT-I TCR. We used this extension of our model to determine if there is a defect in CD8^+^ T cell responses associated with the D2EX mutation and if so, whether this is intrinsic to the T cells, or is a function of other cells, for example the dendritic cells required for priming. To do this, CD8^+^ T cells were purified from the spleens of WT and D2EX OT-I mice also bearing the CD45.1 allelic marker and transferred into groups of WT or D2EX mice (which carry the CD45.2 allele). These mice were infected with HSV.OVA on the next day and after seven days spleens and DRG analysed for the number of OT-I T cells. Irrespective of the recipient genotype, WT OT-I cells expanded in response to infection such that an average of ∼1×10^6^ were found in the spleen. By contrast D2EX OT-I cells failed to expand well with around 10-fold fewer being found (Figure 4A). Likewise, in DRG a significant difference was seen between the numbers of D2EX and WT OT-I at seven days after infection (Figure 4B). In this experiment we also looked at granzyme B (GzmB) expression as a marker of whether the D2EX OT-I might also differ in function, but found that of the OT-I that were recruited to DRG, a similar fraction were Gzm^+^, suggesting adequate differentiation into effectors (Figure 4C). These data suggest that DOCK2 has a significant cell-intrinsic role in ensuring expansion of anti-viral CD8^+^ T cells.

**Figure 4:**
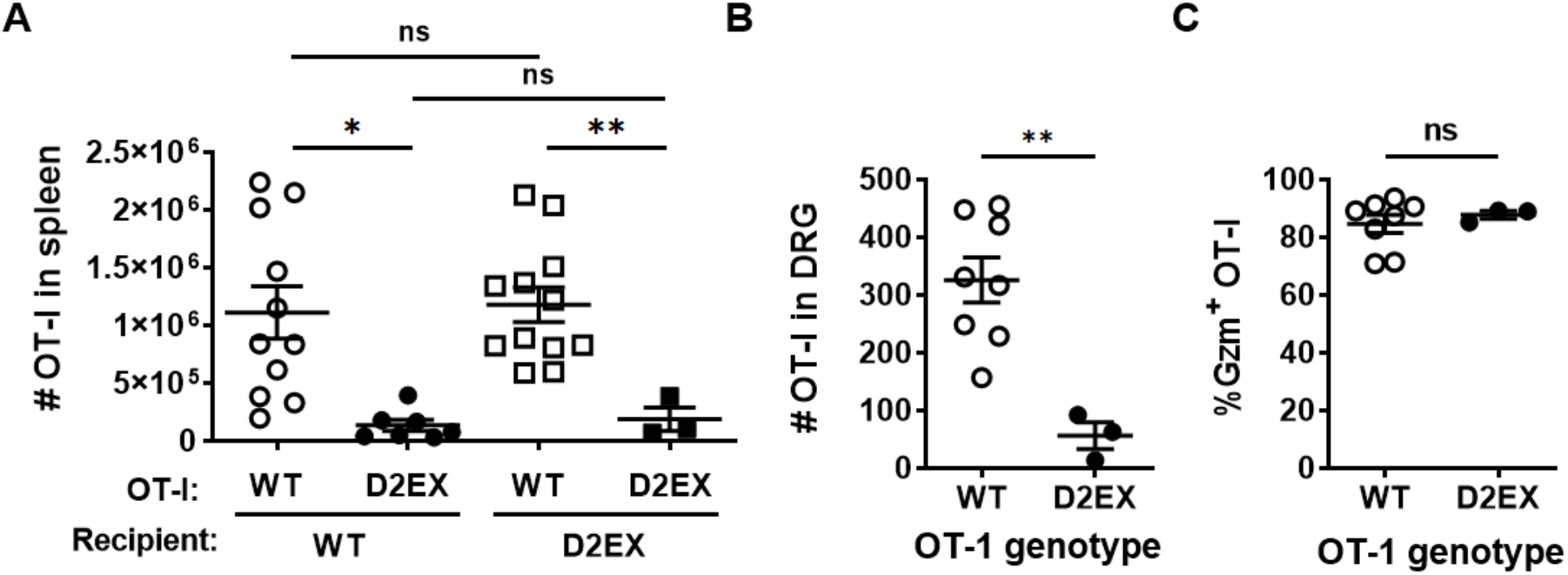
DOCK2 plays a cell-intrinsic role in clonal expansion of anti-viral CD8^+^ T cells. CD8^+^ T cells purified from OT-I mice of the genotypes shown were transferred into WT and D2EX (A) or WT (B, C) mice that were then infected with HSV.OVA 24 hours later. A) Numbers of OT-I T cells in the spleens of mice 7 days after infection, data combined from 3 independent experiments. B) Numbers of OT-I T cells in the DRG, 7 days after infection and the percent of these cells expressing GzmB. Statistical significance was determined using a 2-way ANOVA followed by Sidak’s post-test for pair-wise comparisons (A) or t-tests (B, C); *p<0.05, **p<0.01, ns not significant.

### DOCK2 is required for the full protective effect of anti-viral CD8^+^ T cells

Having found poor expansion of virus-specific D2EX CD8^+^ T cells by HSV infection, but some evidence that differentiation might be unaffected, we wondered next whether any cells that were primed would have anti-viral function. We planned to test this in vivo by first priming and expanding D2EX and WT OT-I T cells in vitro, then transferring these into mice to see how well they might protect against HSV disease. However, first it was necessary to determine whether priming and expansion of D2EX OT-I in vitro was feasible. To do this, WT and D2EX OT-I cells were cultured for 24 hours in the presence of SIINFEKL peptide and examined for initial priming as indicated by upregulation of CD69 as an early activation marker. Surprisingly, there was no difference in CD69 upregulation between WT and D2EX OT-I cells in this experiment, even under limiting peptide stimulation (Fig 5A left). Next we examined IRF4 expression as a marker that indicates the adequacy of priming and predicts clonal expansion [11]. In this case, where almost all WT OT-I strongly upregulated IRF4 by 16 hours and largely maintained this out to 40 hours, this was not the case for D2EX OT-I (Fig 5A right). This provides a likely explanation for the failure of expansion of D2EX OT-I seen in virus infection. We then tried a variety of culture conditions to support enough activation and expansion of D2EX cells to allow transfer into mice and determined empirically that cultures of D2EX OT-I cells supported with 6 ng/ml IL-2, which is twice our usual concentration, grew to similar levels as WT OT-I under standard conditions (3 ng/ml IL-2). Cultures of activated WT and D2EX cells were then transferred into WT mice and a day later they were infected with HSV.OVA on the flank. Activated OT-I cells provided significant protection from lesions caused by HSV.OVA infection irrespective of genotype, suggesting that DOCK2 is not essential for the effector function of anti-viral CD8^+^ T cells (Fig 5B and C and Supplementary Figure 2). However, there was a statistically significant difference in the protection provided by WT and D2EX OT-I cells as determined by peak lesion area, with WT cells being superior. Qualitatively, this meant that those mice that received WT OT-I cells almost all only had small lesions at the inoculation site, without secondary spread to other sites in the dermatome. By contrast mice in the D2EX OT-I group nearly all had some amount of secondary spread. Taken together we conclude that when D2EX CD8^+^ T cells are able to be primed, their anti-viral effector function has a modest defect.

**Figure 5:**
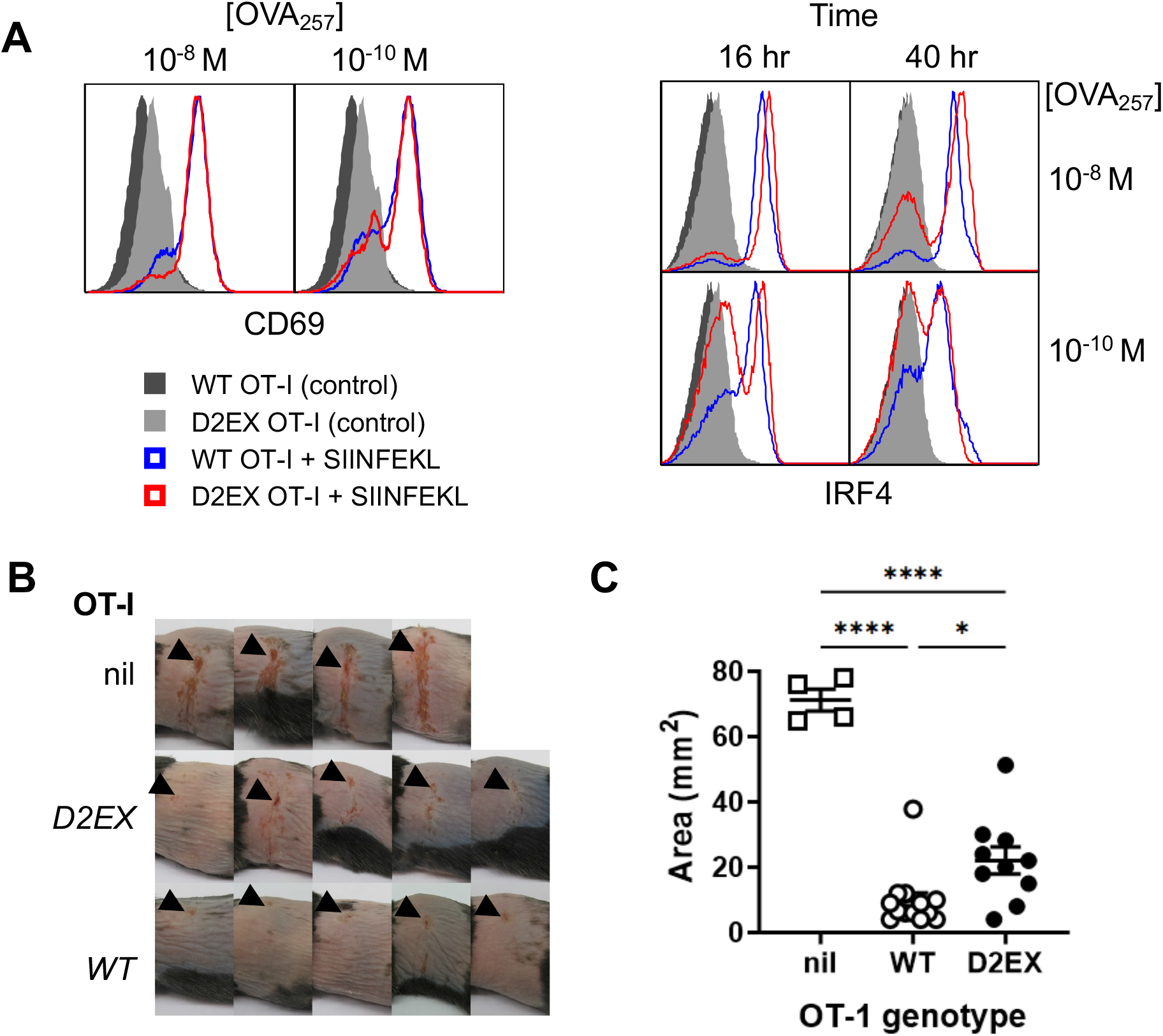
DOCK2 deficent CD8^+^ T cells have slightly reduced protective capacity against HSV infection. CD8^+^ T cells purified from OT-I mice of the genotypes shown were activated with peptide in vitro. A) CD69 and IRF4 were measured at 24, and 16 and 40 hours respectively. B,C) OT-I CD8^+^ T cells were primed and expanded for 4 days and then transferred into WT mice that were infected with HSV.OVA 24 hours later. Images of lesions on mice (B) and peak lesion areas (C) are shown compared with mice that received no cells (nil). Data were combined from two independent experiments; points represent individual mice with bars showing mean and SEM. Statistical significance was determined by 1-way ANOVA with Sidak’s post-test for pair-wise comparisons; ****p<0.0001,*p<0.05, ns not significant.

### Reduced endogenous virus-specific CD8^+^ T cells in D2EX mice

The experiments to date utilised TCR transgenic T cells of a single specificity. To test whether natural CD8^+^ T cell responses to HSV might be affected similarly in D2EX mice, we examined these in the spleen seven days after infection. Just as in unifected mice, the percent and total number of CD8+ T cells were lower in D2EX mice than in WT controls (Figure 6A). CD8^+^ T cells with a TCR specific for the dominant epitope of HSV (gB_498_; SSIEFARL) were also lower in D2EX than WT mice in both analyses and fewer of these cells were expressing GzmB (Figure 6B,C). Finally, the percent and total number of CD8^+^ T cells able to make IFN_γ_ in response to stimulation with SSIEFARL peptide was also reduced in D2EX, compared with WT mice. Taken together, findings from an analysis of the endognous CD8+ T cell response are largely consistent with those gained with transferred OT-I T cells. However, the difference in the size of the response was smaller in the endogenous response and the functional defect, as reflected by Gzm expression and IFN_γ_ production appeared to be more substantial than in the OT-I T cells.

**Figure 6:**
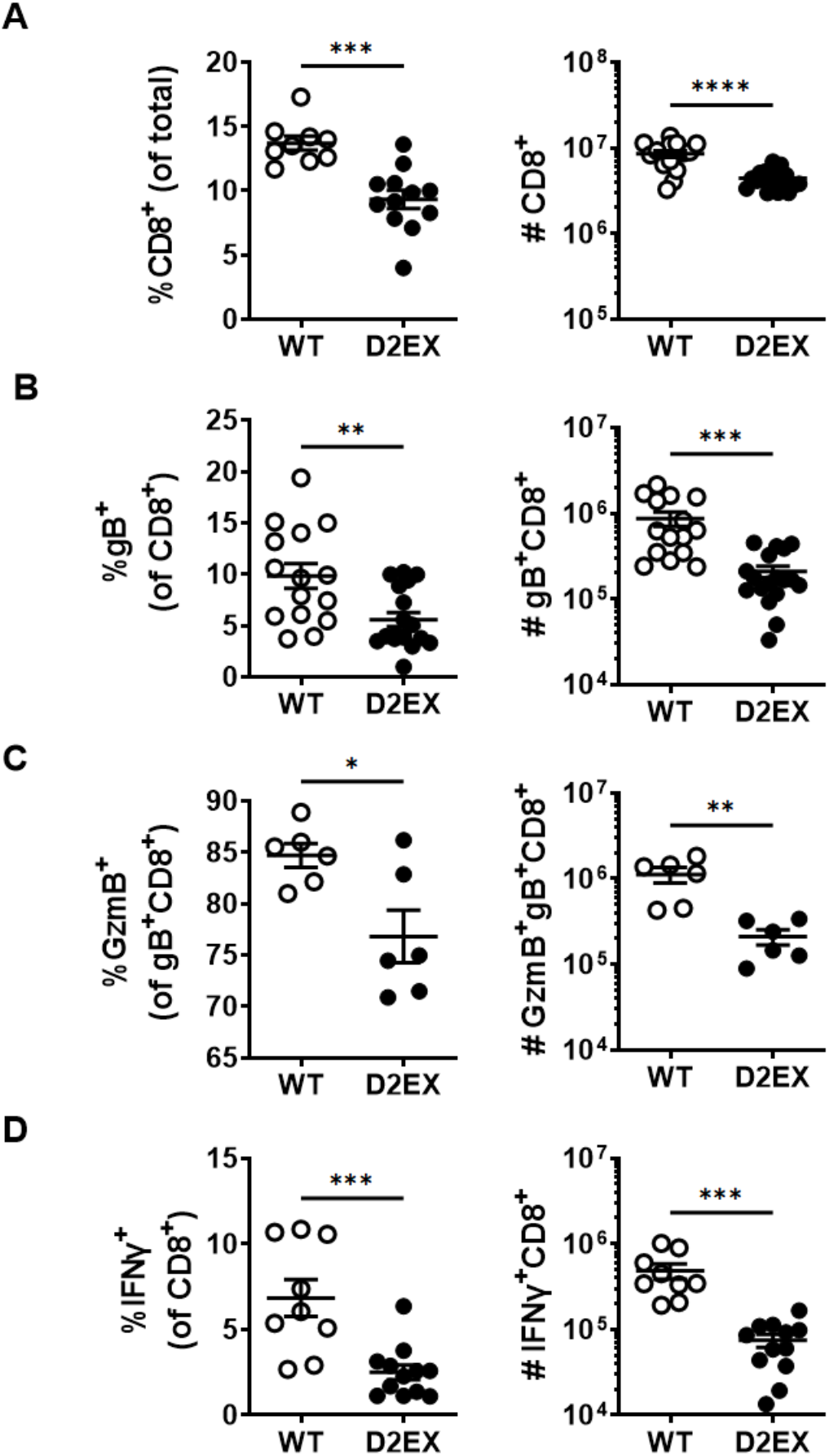
CD8^+^ T cell responses to HSV are deficient in the absence of DOCK2. Mice were infected with HSV and various attributes of CD8+ T cells were measured in spleens 7 days later. Graphs on the left and right of each panel show the percents and total numbers of the populations shown, respectively. A) All CD8+ T cells, B) HSV-gB_498_-specific CD8+ T cells, C) GzmB+, gB_498_-specific CD8^+^ T cells and D) CD8+ T cells able to make IFN_γ_ after stimulation with gB_498_ peptide. Data shown were combined from 5 (A, B), 3 (D) and 2 (C) independent experiments. Statistical significance was determined by t-tests; *p<0.05, **p<0.01, ***p<0.001, ****p<0.0001.

## Discussion

The effect of DOCK2 mutation on mice was first described in 2001 [7]. DOCK2 knock out mice were found to have severe lymphopenia and a chemotactic defect in lymphocytes. Our novel DOCK2 mouse strains recapitulate the previously published phenotypes with absent marginal zone B cells [7], low numbers of NK T cells [18] and T cell lymphopenia [7]. Our characterization studies have also confirmed the previously described apparent “activation” of DOCK2 defective T cells with increased CD44 expression [28], likely due to a peripheral expansion to fill a niche, but in addition, we have shown that this effect can be partially overcome by limiting the T cell repertoire using the transgenic OT-I system. We also show that DOCK2 mice have eosinophilia and elevated levels of IgE on a C57BL6 background, whereas previously this was only shown in TH2 prone Balb/c mice [29].

DOCK2 deficient patients have an increased susceptibility to herpes viruses (particularly CMV and VZV) and this has been ascribed to defects in either T cells or NK cells without a further elucidation which cell type was predominantly responsible for the phenotype [30] as the effect was studied in ex-vivo peripheral blood mononuclear cells (PBMC) from these patients. Using our novel DOCK2 mouse model, we have investigated the role of DOCK2 in the control of herpes virus infections, as these infections are common in DOCK2 deficient patients. Using the HSV mouse model of herpes infection, we show that DOCK2 is important in T cells for control of HSV1 with greater weight loss and higher viral titres in mice lacking DOCK2. We also found that there is a T cell intrinsic defect in priming and expansion of virus-specific CD8^+^ T cells, confirming the importance of DOCK2 in T cells for the control of viral infections. Interestingly, initial in vitro activation of the mutant T cells was normal despite the previously found defect in synapse assembly [5], but the magnitude of expansion of the virus specific CD8^+^ T cells was reduced. We also show that while the cells have anti-viral activity this is also less than in wild type cells including reduced production of interferon-γ by antigen-specific CD8^+^ T cells. This is in agreement with previously published results from patients showing decreased production of interferons by PBMC after 24 hours exposure to HSV1 and vesicular stomatitis virus (VSV) [2]. This work highlights the importance of an infection-based mouse model to investigate the effects of primary immunodeficiencies.

DOCK2 mice have been exposed to *Citrobacter rodentium* previously in an experimental model and show clear defects in innate immunity with increased susceptibility to colitis, more bacterial adhesion and decreased macrophage migration due to the effect of DOCK2 mutations on expression of cytokine receptors [9, 10]. While the HSV model can also highlight defects in innate immunity, the clear defect which we saw was in CD8^+^ T cell antiviral immunity.

We also show here that DOCK2 deficiency results in dysregulation of surface expression of important markers in T cell activation and receptor signaling, with an increased expression of CD5 on both CD4^+^ and CD8^+^ T cells in the absence of infection, indicating an increased TCR signal. As a corollary, we are also the first to describe the “sparing” of FoxP3^+^ cells within the CD4^+^ T cell compartment as these are present at a higher proportion than other T cells subsets in the presence of the severe lymphopenia, with increased TCR signal strength thought to favor the production of Tregs [31]. The only previous literature about regulatory T cells in DOCK2 deficiency showed that co-culture of WT T cells with so called “graft facilitating” cells (defined as CD8^+^ and TCR-cells) isolated from the bone marrow of *Dock2^-/-^* mice failed to induce the formation of FoxP3^+^ or IL10^+^ regulatory T cells [32]. By contrast, here we show that FoxP3^+^ T cells are present in relatively higher numbers in vivo, suggesting that the previous report might have been a result of the used in vitro culture system.

In summary, we show here using a herpesvirus infection model in mice that DOCK2 deficiency leads to defects in T cell immunity, primarily in expansion of cells after priming, but with effector function also compromised. This defect is associated with delayed clearance of infectious virus and prolonged disease and suggests a mechanism for poor control of this family of viruses in people with this immunodeficiency.

## Funding

This study was funded by NIH U19-AI100627; NHMRC through Project Grant 1144684 to AE and KLR and Fellowship 1104329 and Investigator Grants 2008990 to DCT; ACT Private Practice Major Grant 2015 and 2016 to KLR.

### Acknowledgments

We thank the National Computational Infrastructure (Australia) for continued access to significant computation resources and technical expertise; the staff of the Australian Phenomics Facility for animal husbandry; the staff of Australian Cancer Research Foundation Biomolecular Resource Facility for DNA preparation, genotyping, exome and Sanger sequencing; and the staff of the Microscopy and Cytometry Resource Facility for help with flow cytometry. CD1d tetramers used in this study were provided by the NIH tetramer facility (Bethesda, DC).

## Notes

### Competing Interest Statement

The authors have declared no competing interest.

