## Supplementary Figure 1 for "DOCK2-deficiency causes defects in anti-viral T cell responses and poor control of herpes simplex virus infection"

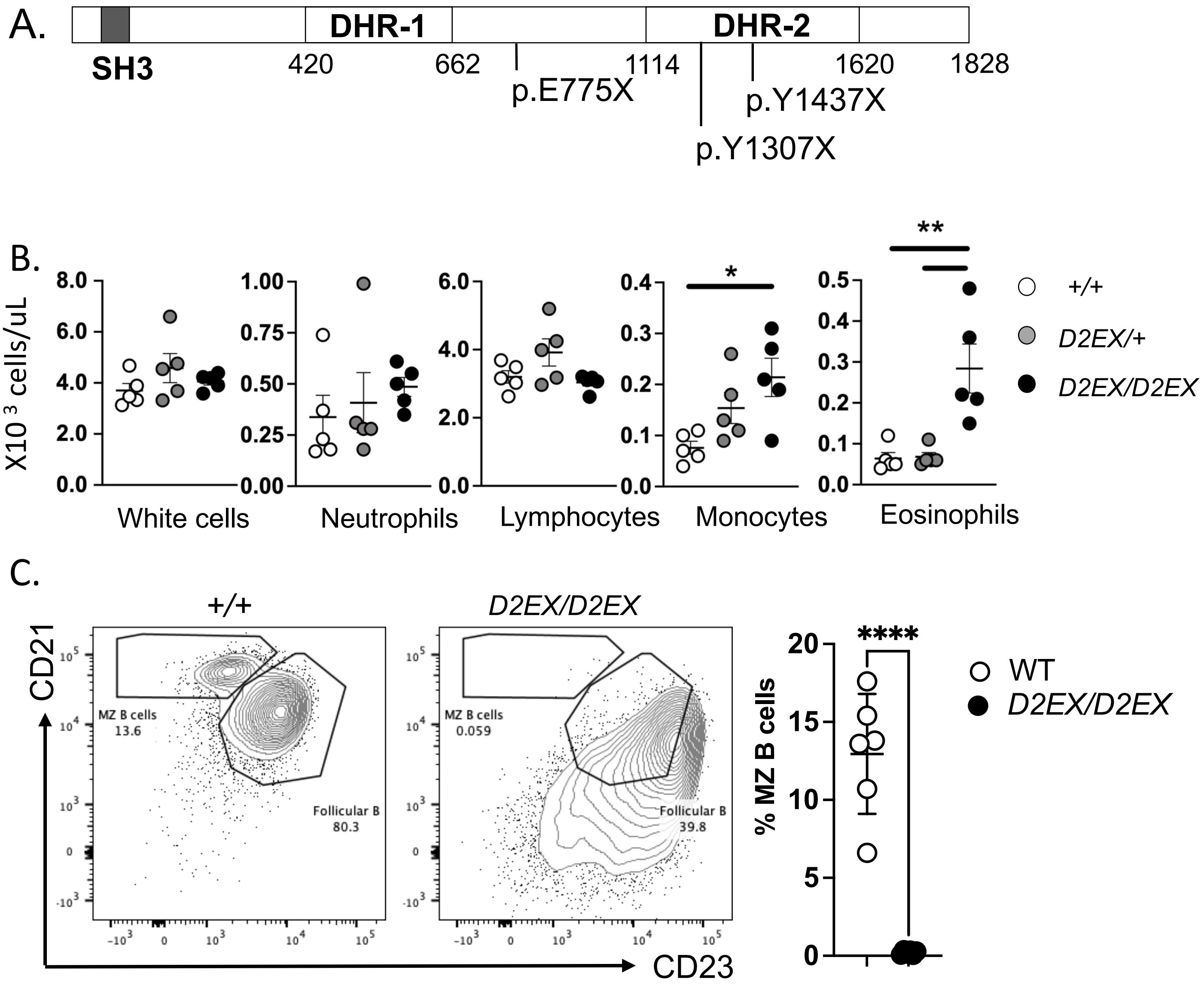


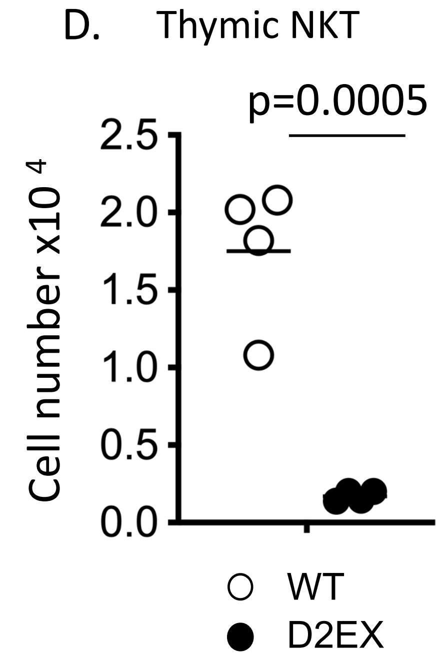


**Supplementary Figure 1**:

A. Graphical representation of the DOCK2 protein structure indicating the sites of the ENU-induced mutations
B. Quantification of the white cell count and differential in the peripheral blood for wild type, heterozygous and mutant mice. Statistical analysis by one way ANOVA with Tukey's multiple comparison test. *<0.05, **<0.005
C. Representative flow cytometry plots (left) and quantitation (right) of marginal zone B cells in the spleen of wild type (white) and mutant (black) mice. Statistical analysis was done by unpaired t-test. ****p<0.0001.
D. Number of thymic NKT cells. Dots represent individual mice, and the mean is indicated by the bar. Statistical analysis by unpaired t –test.
