## Supplementary Figure 2 for "DOCK2-deficiency causes defects in anti-viral T cell responses and poor control of herpes simplex virus infection"

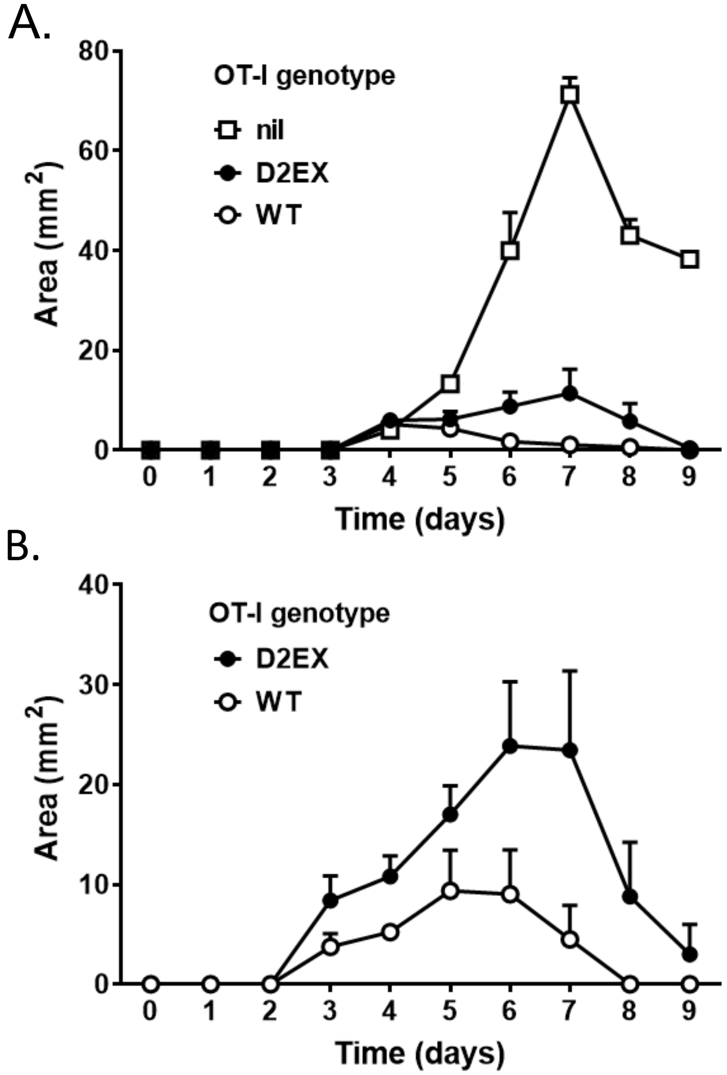


**Supplementary Figure 2. Progression of lesions following HSV infection**.
Progression of lesions in B6 mice that $5x10^6$ received OT-I T cells of the indicated genotype 24 hr before tattoo-infection with HSV.OVA pC-GIP. 2 separate experiments with 4-7 mice per group and experiment.
